# Estimating multiplicity of infection, haplotype frequencies, and linkage disequilibria from multi-allelic markers for molecular disease surveillance

**DOI:** 10.1101/2023.08.29.555251

**Authors:** Henri Christian Junior Tsoungui Obama, Kristan Alexander Schneider

## Abstract

Molecular/genetic methods are becoming increasingly important for surveillance of diseases like malaria. Such methods allow to monitor routes of disease transmission or the origin and spread of variants associated with drug resistance. A confounding factor in molecular disease surveillance is the presence of multiple distinct variants in the same infection (multiplicity of infection – MOI), which leads to ambiguity when reconstructing which pathogenic variants are present in an infection. Heuristic approaches often ignore ambiguous infections, which leads to biased results. To avoid such bias, we introduce a statistical framework to estimate haplotype frequencies alongside MOI from a pair of multi-allelic molecular markers. Estimates are based on maximum-likelihood using the expectation-maximization (EM)-algorithm. The estimates can be used as plug-ins to construct pairwise linkage disequilibrium (LD) maps. The finite-sample properties of the proposed method are studied by systematic numerical simulations. These reveal that the EM-algorithm is a numerically stable method in our case and that the proposed method is accurate (little bias) and precise (small variance) for a reasonable sample size. In fact, the results suggest that the estimator is asymptotically unbiased. Furthermore, the method is appropriate to estimate LD (by *D′, r*^2^, *Q*^*\**^, or conditional asymmetric LD). Furthermore, as an illustration, we apply the new method to a previously-published dataset from Cameroon concerning sulfadoxine-pyrimethamine (SP) resistance. The results are in accordance with the SP drug pressure at the time and the observed spread of resistance in the country, yielding further evidence for the adequacy of the proposed method. The method is particularly useful for deriving LD maps from data with many ambiguous observations due to MOI. Importantly, the method per se is not restricted to malaria, but applicable to any disease with a similar transmission pattern. The method and several extensions are implemented in an easy-to-use R script.

**Author summary:** Advances in genetics render molecular disease surveillance increasingly popular. Unlike traditional incidence-based epidemiological data, genetic information provides fine-grained resolution, which allows monitoring and reconstructing routes of transmission, the spread of drug resistance, etc. Molecular surveillance is particularly popular in highly relevant diseases such as malaria. The presence of multiple distinct pathogenic variants within one infection, i.e., multiplicity of infection (MOI), is a confounding factor hampering the analysis of molecular data in the context of disease surveillance. Namely, due to MOI ambiguity concerning the pathogenic variants being present in mixed-clone infections arise. These are often disregarded by heuristic approaches to molecular disease surveillance and lead to biased results. To avoid such bias we introduce a method to estimate the distribution of MOI and frequencies of pathogenic variants based on a concise probabilistic model. The method is designed for two multi-allelic genetic markers, which is the appropriate genetic architecture to derive pairwise linkage-disequilibrium maps, which are informative on population structure or evolutionary processes, such as the spread of drug resistance. We validate the appropriateness of our method by numerical simulations and apply it to a malaria dataset from Cameroon, concerning sulfadoxine-pyrimethamine resistance, the drug used for intermittent preventive treatment during pregnancy.

## Introduction

Genomic/molecular techniques allow the detection of pathogens with better granularity than the traditional clinical methods and, therefore, enhance disease surveillance [1, 2]. Molecular surveillance of infectious diseases allows for retrieving and reconstructing population or evolutionary genetic information such as the spread of drug-resistant pathogenic variants, or detailed epidemiological information such as routes of transmission and transmission intensities [3]. Such information helps to optimize disease control and prevention and becomes increasingly popular for some of the economically most relevant infectious diseases, such as malaria [4, 5].

A number of programs targeting the reduction of the global malaria burden have been launched in the past decade, e.g., the President’s Malaria Initiative (PMI), the UN Millennium Development Goals (MDGs) followed by the UN Sustainable Development Goals (SDGs; Goal 3 Good Health and Well-Being) accompanied by the Global Technical Strategy For Malaria [6, 7]. Despite substantial reductions in malaria incidence in the past decades, the trend has been reversed since 2018. With an estimated 247 million infections, and, 621 000 deaths worldwide in 2022 the disease remains highly endemic, particularly in sub-Saharan Africa, and parts of Oceania (i.e., Papua New Guinea), and South America [8]. This challenges the goal of malaria elimination in at least 30 countries by 2030.

Challenges in successful malaria control are, e.g., the spread of drug resistance, or migration events, by which the disease is reintroduced into areas aiming for malaria elimination and causes local disease outbreaks [9, 10]. Both, the spread of drug resistance and migration events cause characteristic patterns of linkage-disequilibrium (LD) which are informative about the underlying evolutionary-genetic processes [11]. This emphasizes the importance of harnessing LD measures in malaria molecular surveillance.

Pairwise LD maps require reliable haplotype frequency estimates. Such estimates are not straightforward in malaria, because multiple genetically distinct pathogenic variants can be present within the same infection, often referred to as complexity of infection (COI) or multiplicity of infection (MOI) [12–14]. Because of this confounding factor, *Plasmodium* (although a haploid organism) ‘behaves’ as a polyploid organism with a random level of ploidy within human infections.

The characteristic malaria transmission cycle in combination with MOI leads to a departure from classical population genetics [15, 16]. Notably, malaria transmission involves a step of meiotic recombination in the mosquito vector, implying that *Plasmodium* acts as a diploid organism within the mosquito. Note that the terms COI or MOI are defined ambiguously in the literature. Here, we follow the stringent definition of [17] of MOI as the number of super-infections, i.e., the number of infective events with the same or different pathogenic variants during one disease episode.

Therefore, care has to be taken when estimating the population genetics of such diseases, as it differs from classical population genetics due to MOI. For instance, MOI plays an important role in the estimation of the frequency of pathogenic variants, and more importantly in the differentiation between pathogens frequencies and prevalence [17].

Typically, statistical models are required to estimate the frequency of each haplotype in the pathogen population from molecular data, while accounting for MOI. Note that, in low transmission areas, only a relatively small amount of infections occurs, yielding molecular dataset with small sample sizes. Moreover, molecular data usually contains missing values as a result of failures in molecular assays and errors in the determination of alleles at makers of concern [17]. The amount of missing values per sample increases with the number of molecular markers considered. Therefore, to minimize the loss of data, fewer loci can be considered. For studying pairwise LD, it suffices to consider two marker loci.

Several statistical methods have been developed to estimate haplotype frequencies alongside MOI in malaria from molecular data. The maximum-likelihood (ML)-based method of [18] estimates haplotype frequencies and MOI, considering one polymorphic maker locus. Another maximum-likelihood method, developed by [19] estimates haplotype frequencies alongside MOI from multiple bi-allelic SNPs and presents a framework to estimate haplotype prevalence using the frequency and MOI estimates as plugin parameters. The method is of particular interest if applied to estimate the frequency of drug-resistant haplotypes in malaria, which are typically characterized by five to ten SNPs [20]. Although this method allows monitoring the spread of drug resistance, mining the patterns of selection caused by drug-resistance evolution (genetic hitchhiking, LD) requires different molecular data and analysis methods. Variable molecular markers such as STRs are useful in this regard, as they are fast evolving and allow to investigate ongoing evolutionary processes such as drug resistance evolution. To study genetic hitchhiking in terms of heterozygosity, it is sufficient to estimate allele-frequency spectra separately for a set of markers flanking the loci under selection, e.g., by the method of [18]. To study pairwise LD maps, haplotypes determined by two multi-allelic markers have to be considered.

For this purpose, we introduce a statistical model that allows to estimate haplotypes frequencies and MOI by ML, assuming two multi-allelic loci. Because no explicit solution for the maximum-likelihood estimate (MLE) can be found, we employ the expectation maximization (EM)-algorithm (cf. [19, 21]) to obtain a numerically efficient and stable method to derive the estimates. A systematic simulation study is performed, to evaluate the finite sample properties of the MLE. Furthermore, we investigate the capability of the model to estimate the multi-allelic LD measures *D′, Q*^*\**^, and *r*^2^ (cf. [22]).

As an example, the method is applied to a molecular dataset from *P. falciparum*-positive (*n* = 166) blood samples collected in Yaound’e, Cameroon from 2001-2002 and 2004-2005 [11]. The patterns of pairwise LD around the dhfr and dhps genes, responsible for sulfadoxine-pyrimethamine (SP) resistance, are studied in terms of mulltiallelic LD measures including the conditional asymmetric LD measure proposed by [23].

An implementation of the maximum-likelihood method is provided as an easy-to-use R script accessible via GitHub https://github.com/Maths-against-Malaria/MultiAllelicBiLociModel and on Zenodo at https://doi.org/10.5281/zenodo.8289710.

The statistical model is described in Methods, while the model’s performance and data applications are described in Results. Readers primarily interested in the applications shall go straight to Results. However, readers interested in the mathematical details, find a concise description of all derivations in S1 Mathematical appendix.

## Results

Estimating pathogen haplotype frequencies is a fundamental basic to monitor, for instance, haplotypes of interest (e.g. drug-resistance-associated haplotypes) or to reconstruct past or ongoing evolutionary events such as selection. The latter is typically investigated by considering summary statistics such as pairwise linkage disequilibria (LD) [22, 24], which are calculated from haplotype frequencies.

Assuming a genetic architecture of two multi-allelic loci, we first show how the MLE is derived. Then we provide an application to empirical data and contrast different LD measures. Finally, we investigate the finite sample properties of the estimator.

### The maximum-likelihood estimate

The estimates for haplotypes frequencies and MOI are obtained by maximizing the likelihood function (12). Because the likelihood function (12) is high-dimensional and non-linear, no closed solution could be found. Therefore, the likelihood function has to be maximized numerically. Applying Newton or Quasi-Newton methods to maximize the likelihood function over a multi-dimensional space (particularly over the simplex as in our case) is inconvenient in practice because convergence is too sensitive to initial conditions. Therefore, we pursue with the expectation maximization (EM)-algorithm. The algorithm is typically insensitive to initial conditions but tends to be slow close to the point of convergence (cf. [25] Chapter 2).

The EM-algorithm is a two-step recursive method consisting of (i) the expectation-step (E) and (ii) the maximization-step (M). Starting from an initial parameter choice ***θ***_0_, it updates the parameter vector ***θ***_*t*_ in every step *t* until convergence is reached numerically, to yield the MLE. First the log-likelihood of the unobserved MOI, i.e., (*m*, ***m***) which are random variables, and the unknown parameters ***θ***, given the observed data is calculated. At iteration *t*, in the E-step, the expectation of this likelihood function is calculated with respect to the distribution of the unobserved random variable, given the observed data and the parameter choice ***θ***_*t*_. In the M-step the resulting function is maximized with respect to the unknown parameters ***θ***, which yields the updated parameters vector ***θ***_*t*+1_. We derive the algorithm in section Maximum-likelihood estimate of the S1 Mathematical appendix in detail.

In the present case, the EM-algorithm yields the following iteration. First, arbitrary initial values of the MOI (Poisson) parameter *λ*^(0)^ and haplotype frequencies ***p***^(0)^ are chosen. First, in step *t* + 1, the frequency estimate of each haplotype ***h*** is updated as

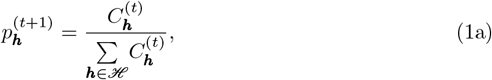

where

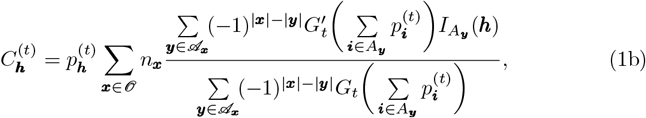

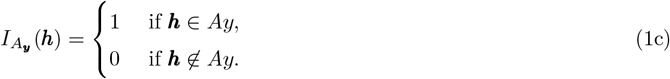

and *G*_*t*_(*z*) is the PGF (3b) with parameters in step *t*. (Note that eq. 1 is not restricted to the conditional Poisson distribution.)

At the second step of the algorithm, the MOI parameter *λ*_*t*+1_ is updated by iterating the equation

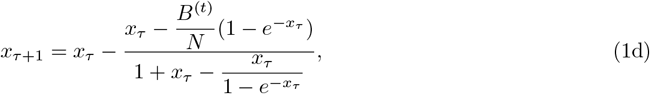

where

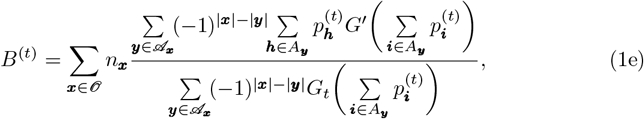

starting from *x*_0_ = *λ*_*t*_ until convergence is reached numerically. Convergence is reached if |*x*_*τ*+1_ *− x*_*τ*_ | *< ε* holds. The MOI parameter is updated as *λ*_*t*+1_ = *x*_*τ*+1_.

These two steps are repeated until convergence is reached numerically. More precisely, the algorithm terminates at step *t* + 1 if 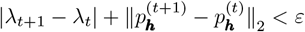 holds. The MLE is then given by

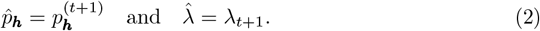

Depending on the application, the haplotype-frequency estimation is of primary interest, while there is prior information concerning MOI. The EM-algorithm can be easily modified to allow the possibility of inputting a plugin estimate of the MOI parameter. The EM-algorithm can be adapted to this case following [19]. This is described in section The EM-algorithm using a plugin estimate for the MOI parameter in S1 Mathematical appendix.

The EM-algorithm (with and without a plugin estimate for the MOI parameter) is implemented as an R script available in the supplementary material, at https://github.com/Maths-against-Malaria/MultiAllelicBiLociModel and on Zenodo at https://doi.org/10.5281/zenodo.8289710. Importantly, the implementation also allows deriving bootstrap-percentile confidence intervals (CIs) and bootstrap bias corrections of the estimates (cf. [26]) for the MLEs. A detailed description of the usage of this script is provided in S1 User Manual.

### Data application

We estimate linkage disequilibrium (LD) from molecular data of malaria parasites extracted from blood samples, which were positive for *P. falciparum*, collected in Yaound’e, Cameroon from 2001 to 2002 (*N* = 166 samples) and 2004 to 2005 (*N* = 165 samples). The dataset was previously described in [11]. The data contains information associated with sulfadoxine-pyrimethamine (SP) resistance [27–29]. Particularly, allelic information is available at (i) codons 51, 59, 108, and 164 at *Pfdhfr*, (ii) codons 436, 437, 540, 581, and 613 at *Pfdhps*, (iii) 18 STR markers on chromosome 4 flanking *Pfdhfr*, (iv) 15 STR markers on chromosome 8 flanking *Pfdhps*, and (v) 8 neutral STR markers at chromosome 2 and 3 [11]. Because no mutations occurred on codon 164 at *Pfdhfr* and codons 540 and 581 at *Pfdhps*, they were excluded from pairwise LD analysis. The remaining six SNPs were all bi-allelic.

Pairwise LD was measured by *D′, r*^2^, *Q*^*\**^, and ALD. We estimated pairwise LD for (i) the 6 SNPs retained at *Pfdhfr* and *Pfdhps*, and (ii) all the 41 STR makers. Reported CIs are bootstrap percentile CIs [26].

Fig. 1 shows pairwise LD measured by *D′* and ALD for the 6 polymorphic SNPs at *Pfdhfr* and *Pfdhps*. In general, LD was high at *Pfdhfr* in the years 2001/2002: *D′* = 0.79 (95% CI: [0.62, 0.95]) between codons 51 and 59, *D′* = 1 (95% CI: [1, 1]) between codons 51 and 108, and *D′* = 0.95 (95% CI: [0.85, 1]) between 59 and 108 (Fig 1A). This is not surprising since pyrimethamine resistance is acquired sequentially, starting with a base mutation at codon 108, which is followed by a secondary mutation either at codon 51 or 59 to cause intermediate resistance [28]. A combination of the three mutations then causes high levels of resistance [28]. The mutations at *Pfdhps* are associated with sulfadoxine resistance [28]. Levels of LD were generally lower (Fig 1A) than at *Pfdhfr*, which coincides with the empirical observations elsewhere (e.g. [20]). High LD values between some SNPs at *Pfdhfr* and *Pfdhps* are indicated by *D′* (Fig 1A) and neither by ALD (Fig 1C) nor *r*^2^ (coinciding with *Q*^*\**^; S1 Fig). This is an artifact of *D′* which occurs for unbalanced frequency distributions, when some haplotypes are absent because of low allele frequencies. The corresponding values of ALD (symmetric in this case) are substantially smaller. The values of *r*^2^ (*Q*^*\**^) are the squared ALD values in this case and hence close to zero.

**Fig 1.**
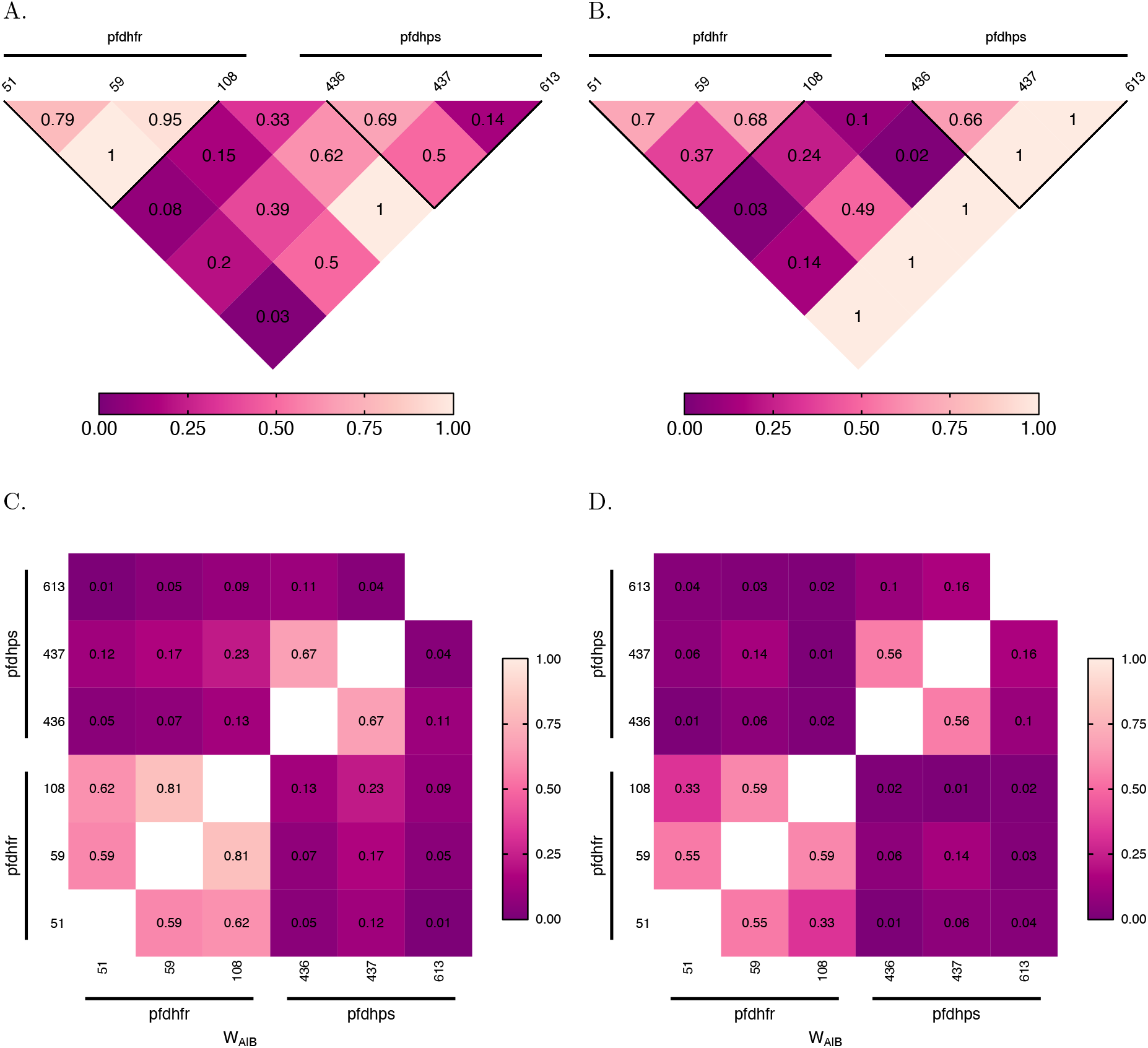
Linkage disequilibrium at SNP markers: Shown are estimates of pairwise LD for six SNP marker loci associated with SP drug-resistance, i.e., three codons (51, 59, 108) at *Pfdhfr* and three codons (436, 437, 613) at *Pfdhps*. Panels (A, B) show *D*^*I*^ values, whereas (C, D) show ALD values. Furthermore, panels (A, C) correspond to the years 2001/2002, while panels (B, D) correspond to the years 2004/2005. The codons on *Pfdhfr* and *Pfdhps* are highlighted on the maps by horizontal black lines, with the name of the gene and codons are specified on top and below the line, respectively. The thick black lines in panels (A, B) group pairwise LD within each gene. The numbers indicate LD values.

Overall, these results are indicative of recent and independent spreads of pyrimethamine (*Pfdhfr*) and sulfadoxine (*Pfdhps*) resistance, which were still ongoing in 2001/2002. The results are in agreement with the high level of SP drug pressure during that time, when chloroquine was discontinued as first-line treatment and replaced by amodiaquine as first-, and SP as second-line treatment [11]. ALD values (Fig 1C) are in agreement with *D′*. Because *r*^2^ and *Q*^*\**^ are the squared ALD measure, they yield values close to zero (S1 Fig).

The LD map for the years 2004/2005 reveals a decrease of pairwise LD between codons at *Pfdhfr*, i.e., *D′* = 0.7 (95% CI: [0.42, 1]) between 51 and 59, *D′* = 0.37 (95% CI: [0.12, 0.67]) between 51 and 108, and *D′* = 0.68 (95% CI: [0.36, 1]) between 59 and 108 (see Fig 1B).

The reason for the decrease is the independent mutational origins at codons 51 and 59 succeeding the mutation at codon 108. While the wildtype and 108 single mutant decreased in frequency, the 51/108 and 59/108 double mutants as well as the 51/59/108 triple mutant increased in 2004/2005. Single mutations at codons 51 and 59 were almost absent in 2001/2002, but the 51/108, 59/108 double mutants, and the 51/59/108 triple mutant were abundant. This led to strong LD in 2001/2002. However, in 2004/2005 the 51 and 59 single mutations were rarely observed. Because of the overall very unbalanced haplotype frequencies, this led to a particularly strong drop in LD between codons 51 and 108. These results have to be considered with caution, as they are likely sampling artifacts in combination with the particularities of LD measures for highly unbalanced distributions. This is reflected by the wide CIs in 2004/2005.

Looking at the LD map for the microsatellite data (Fig 2), similar patterns are observed in the years 2001/2002 and 2004/2005. High LD is observed around the *Pfdhfr* gene (chromosome 4) (Fig 2), indicative of relatively strong selection on *Pfdhfr*, being reflected by the high frequency of resistance-associated mutations (cf. [19], Table 3). LD is also observed around *Pfdhps*, but to a lesser extent, in agreement with the observation that selection is weaker than for *Pfdhfr* ([20]). The reduced variation flanking *Pfdhfr* and *Pfdhps* result in intermediate ALD between the markers flanking *Pfdhfr* (chromosome 4) and chromosome 2 as well as between the markers flanking

**Fig 2.**
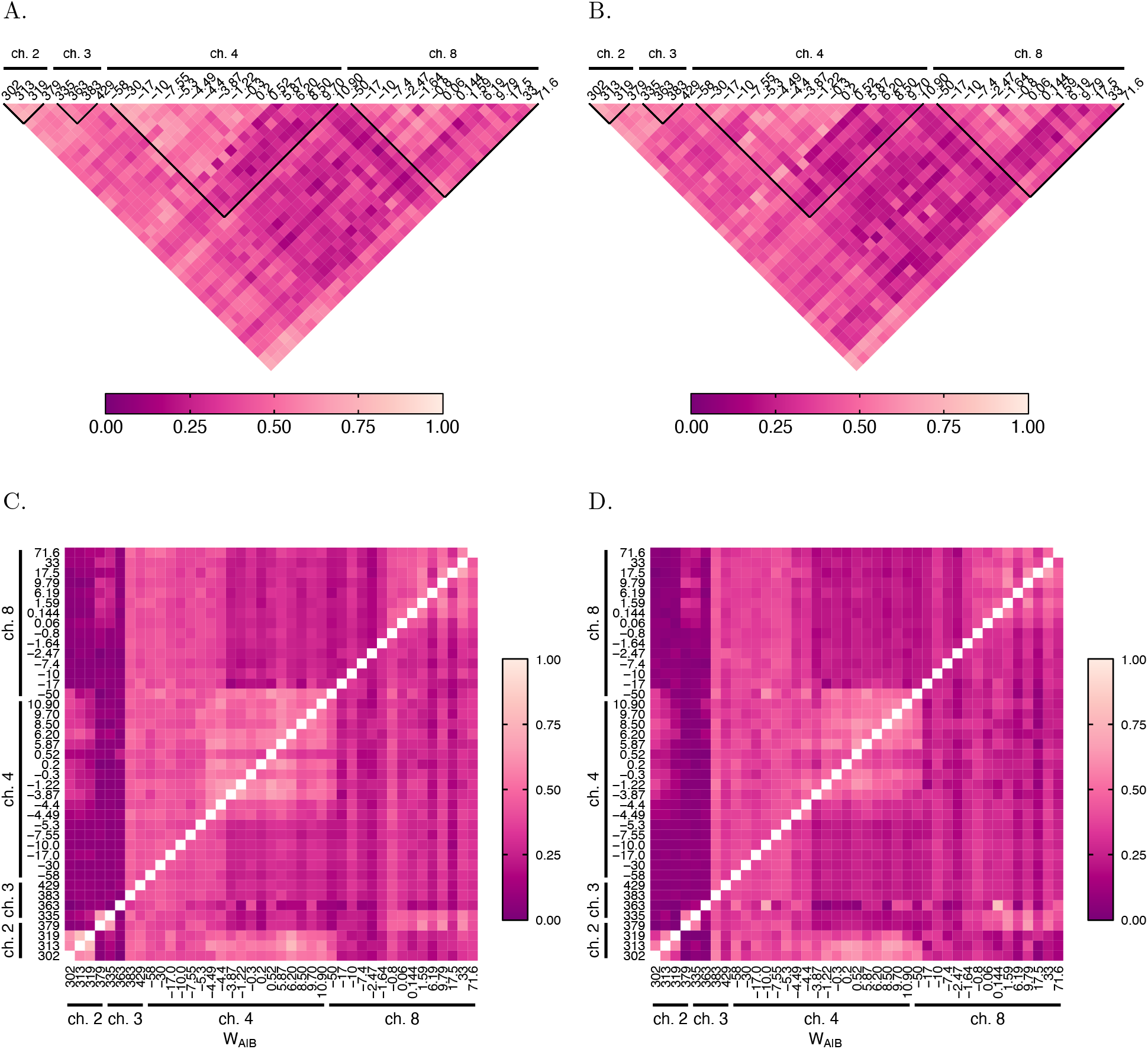
Linkage disequilibrium at microsatellite markers: As in Fig 1 but for a set of genes at two neutral chromosomes, i.e., chromosomes 2 and 3, genes around *Pfdhfr* on chromosome 4, and genes around *Pfdhps* on chromosome 8. *Pfdhps* (chromosome 8) and chromosome 3. The high LD values at chromosomes 2 and 3 are sampling artifacts. First, all markers at chromosomes 2 and 3 are highly polymorphic. Consequently, the number of haplotypes that can be formed by all allelic combinations exceeds sample size by far. Second, at each marker a substantial amount of samples failed to successfully amplify, resulting in large amounts of missing data. I.e., for any pair of markers at chromosome 2 and 3 the number of samples, with missing data at one or both markers is high. This resulted in the artifact that many haplotypes with distinct alleles at both markers were observed, yielding high LD.

### Evaluation of the estimator’s performance

Estimates for the MOI parameter and haplotype frequencies are subsumed by 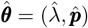 in the following. Maximum-likelihood estimators typically have desirable asymptotic properties (as proven for a simpler case in [30]). However, the performance of the estimator has to be investigated for finite sample size. Ultimately, a good estimator is accurate, i.e., low bias, and precise, i.e., low variance. Here, these two components of performance are investigated by numerical simulations because the MLE has no closed form solution.

Together, bias and variance yield the mean squared error (MSE), defined by 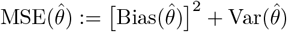. The assessment of an estimator using the MSE implies a trade-off between variance and bias.

#### Bias of the estimator

Here, to facilitate the comparison of bias across different parameter values, we estimate bias as a normalized quantity, namely the relative bias (14a) in percent.

##### Relative bias for haplotypes frequencies estimates

In the case of a balanced true haplotype distribution, the estimator appears to be unbiased, irrespectively of sample size and number of alleles *n*_1_ and *n*_2_ considered at each locus (see Fig 3). This result is expected, because for a balanced distribution, the haplotypes are equally represented in the population, making them equivalent. High values of relative bias in this case, are simulation artifact, explained by random sampling, which effect is emphasized for small sample sizes and large number of alleles *n*_1_ and *n*_2_.

**Fig 3.**
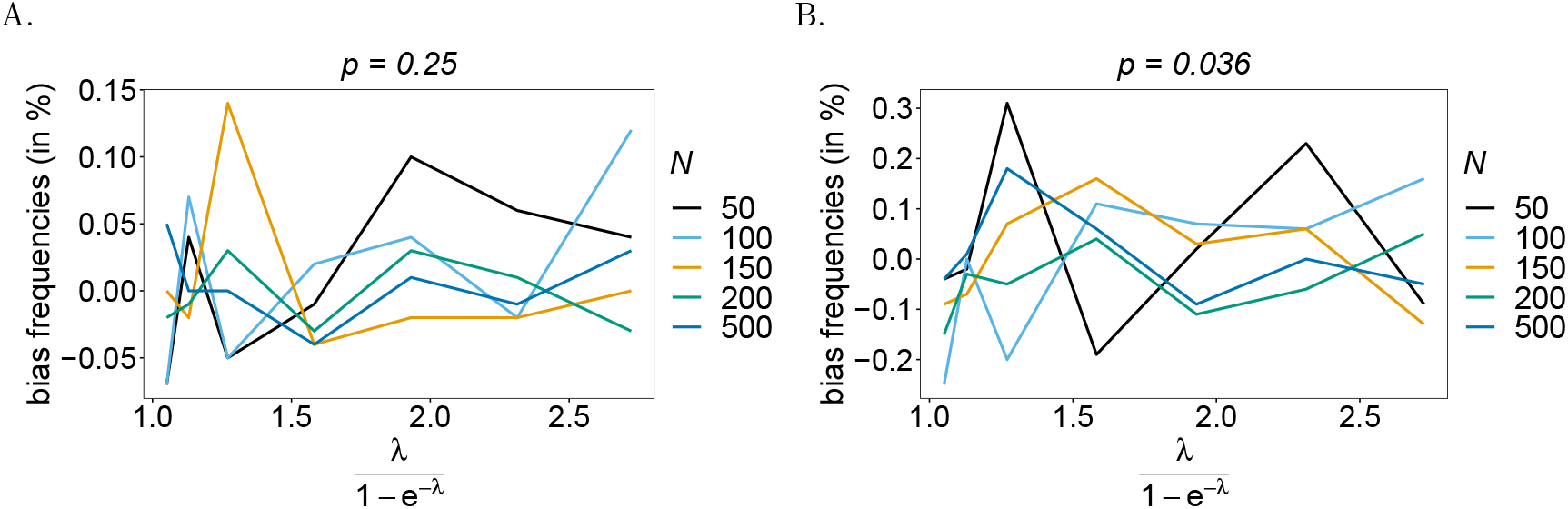
Bias of frequencies estimates – the balanced case: Shown is the bias of the frequency estimates in % of the true value, as a function of the mean MOI (i.e., for a range of MOI parameters). The balanced haplotype frequency distributions given in Table 1 for *n*_1_ = *n*_2_ = 2 (A) and for *n*_1_ = 4, *n*_2_ = 7 (B). In both panels, only the bias for the first haplotype is shown (in the balanced case, all haplotypes are equivalent). Colors correspond to different sample sizes.

Bias remains low for an unbalanced true haplotype frequency distribution (see Fig 4). Haplotypes with low frequency are under-represented in datasets, hence, are estimated with a higher bias than predominant haplotypes that are over-represented. Therefore, haplotype frequencies will be overestimated for predominant haplotypes and underestimated for the under-represented ones. This is particularly noticeable for high mean MOI (*ψ >* 1.8) and small sample size (*N <* 150). A comparison of bias for both the predominant and under-represented haplotypes for *n*_1_ = *n*_2_ = 2 (see Figs 4A and B) with the bias of those haplotypes for *n*_1_ = 4 and *n*_2_ = 7, shows that bias tends to increase with the number of alleles considered. In fact, an increase in *n*_1_ and *n*_2_ yields a geometrical increase in the number of haplotypes. Therefore, assuming the same frequency of predominant haplotypes, the frequencies of some remaining haplotypes have to decrease for larger values of *n*_1_ and *n*_2_.

**Fig 4.**
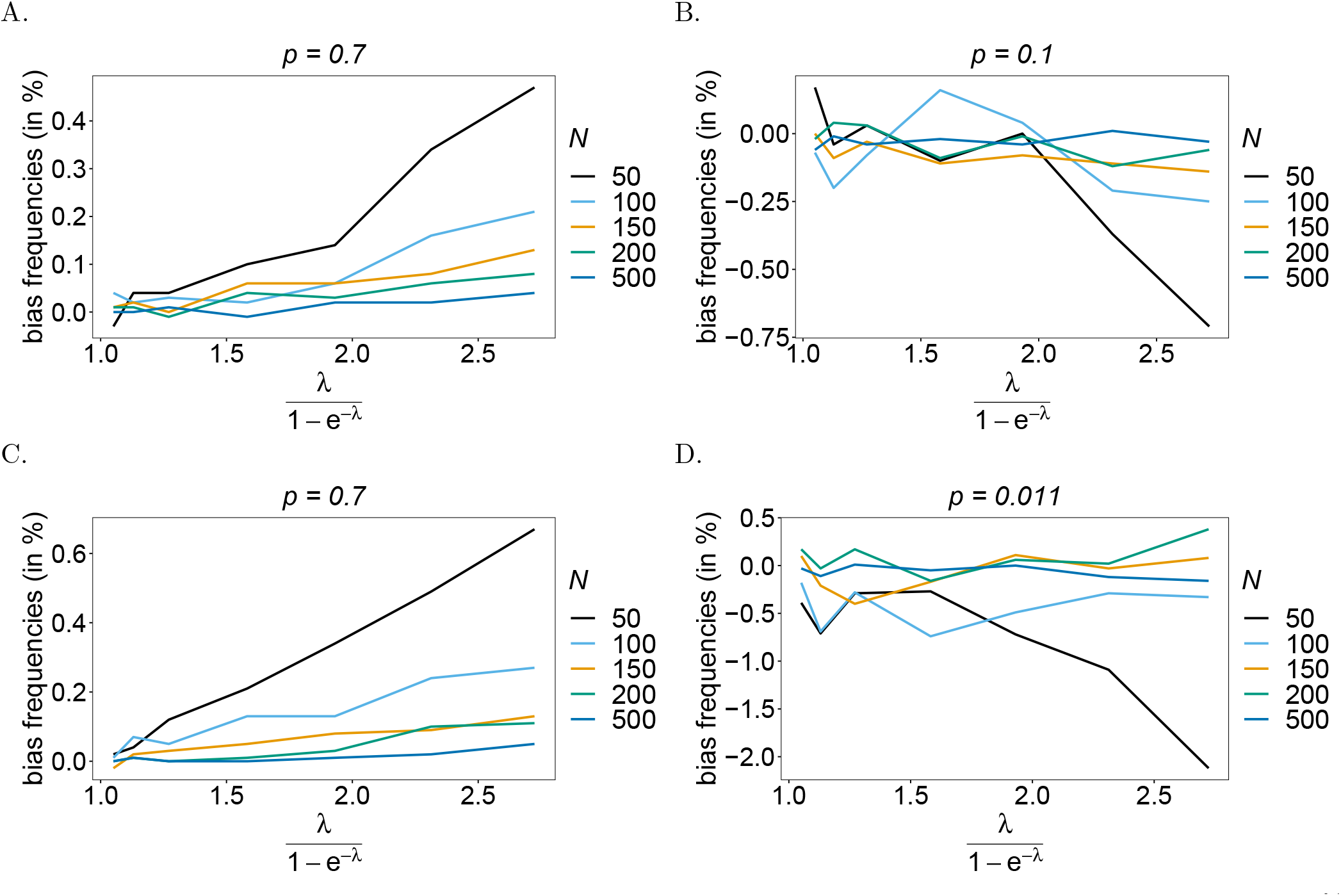
Bias of frequencies estimates – the unbalanced case: Shown is the bias of the frequency estimates in % of the true value, as a function of the mean MOI (i.e., for a range of MOI parameters). The unbalanced haplotype frequency distributions given in Table 1 for *n*_1_ = *n*_2_ = 2 are assumed in (A, B), while those for *n*_1_ = 4, *n*_2_ = 7 are assumed in (C, D). In both cases, only the bias for the predominant haplotype and one underrepresented haplotype are shown (all underrepresented haplotypes are equivalent). Colors correspond to different sample sizes.

##### Relative bias for MOI parameter estimates

Estimates for the mean MOI *ψ* are empirically more relevant than estimates for the MOI parameter *λ*. Therefore, we evaluated the bias for *ψ* instead of *λ*. Overall, the estimator has low bias for estimates of mean MOI (see Fig 5). Note that *λ* and therefore *ψ* have a lower but no upper bound. Hence, they can be overestimated by an unlimited amount, but not arbitrarily underestimated. For larger *n*_1_ and *n*_2_, and small sample sizes, rare super-infections with many different haplotypes are occasionally over-represented, yielding a substantial overestimation of *λ* and *ψ*. Therefore, bias tends to increase with *ψ* and decrease with sample size. For unbalanced true haplotype distributions, rare haplotypes are under-represented in datasets, and – particularly for large *ψ* – single infections with the rare haplotypes are unlikely, which yields a higher bias (see Figs 5C, D).

**Fig 5.**
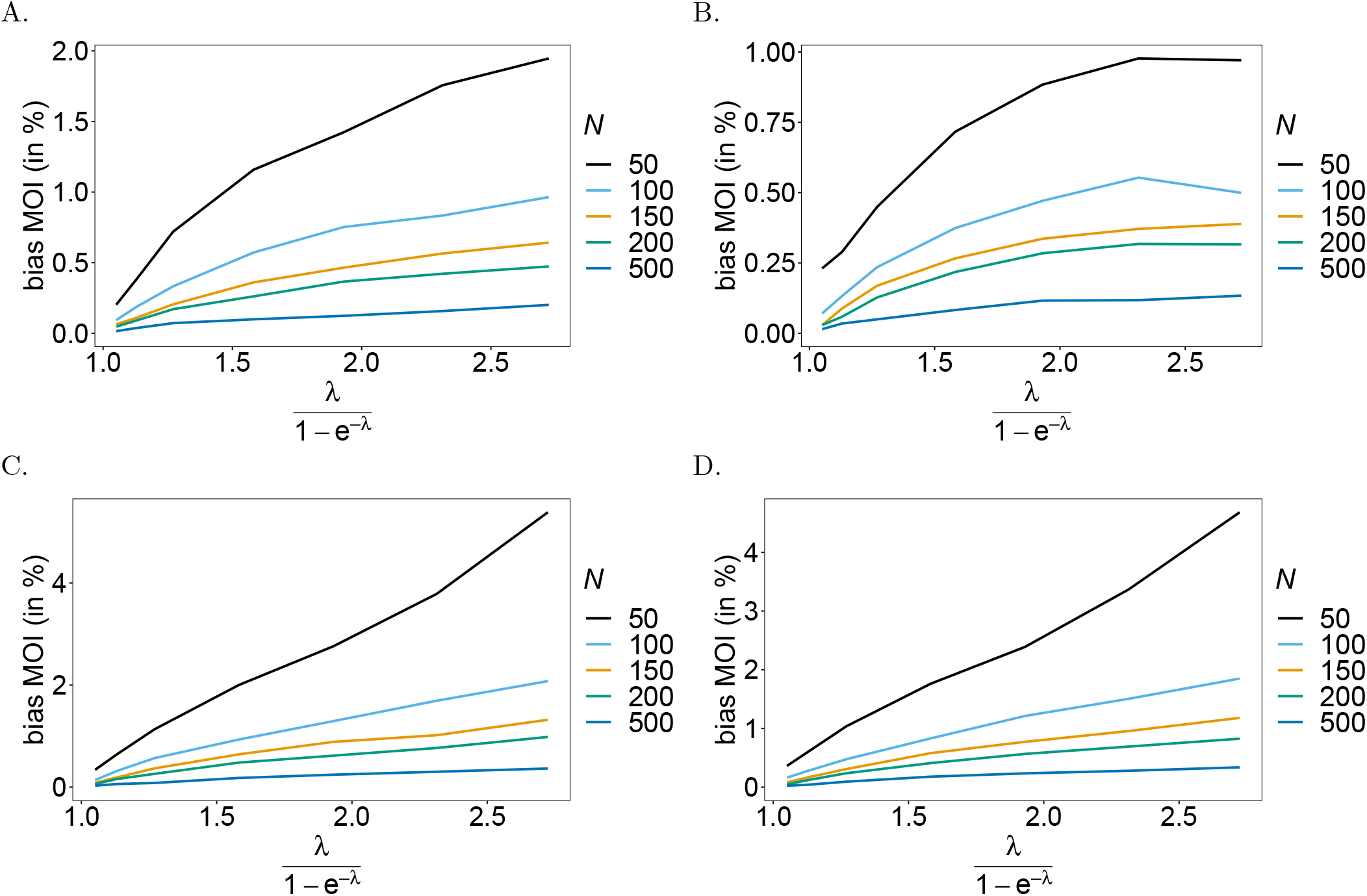
Bias of MOI estimates: Shown is the bias of the mean MOI estimates in % as a function of the true mean MOI (i.e., for a range of MOI parameters). Balanced haplotype frequency distributions (Table 1) are assumed for *n*_1_ = *n*_2_ = 2 (A) and *n*_1_ = 4, *n*_2_ = 7 (B), whereas unbalanced distributions are assumed in panels (C) and (D), for *n*_1_ = *n*_2_ = 2 and *n*_1_ = 4, *n*_2_ = 7, respectively. Colors correspond to different sample sizes.

#### Variance of the estimator

The variance is only reported for the mean MOI. The estimator’s variance is evaluated across a range of parameter values, which requires to normalize variance for better comparison. Hence, the estimator’s variance was assessed using the coefficient of variation (14b).

In general, the variance of the estimator is small. A comparison between the case *n*_1_ = *n*_2_ = 2 and *n*_1_ = 4, *n*_2_ = 7 shows that variance increases with the number of alleles present at each locus (compare Figs 6A, B and C, D). Due to the geometrical increase of the number of haplotypes as *n*_1_ and *n*_2_ increase, the true underlying population is less accurately represented. As expected, variance decreases with sample size as the dataset is more representative of the true underlying population.

**Fig 6.**
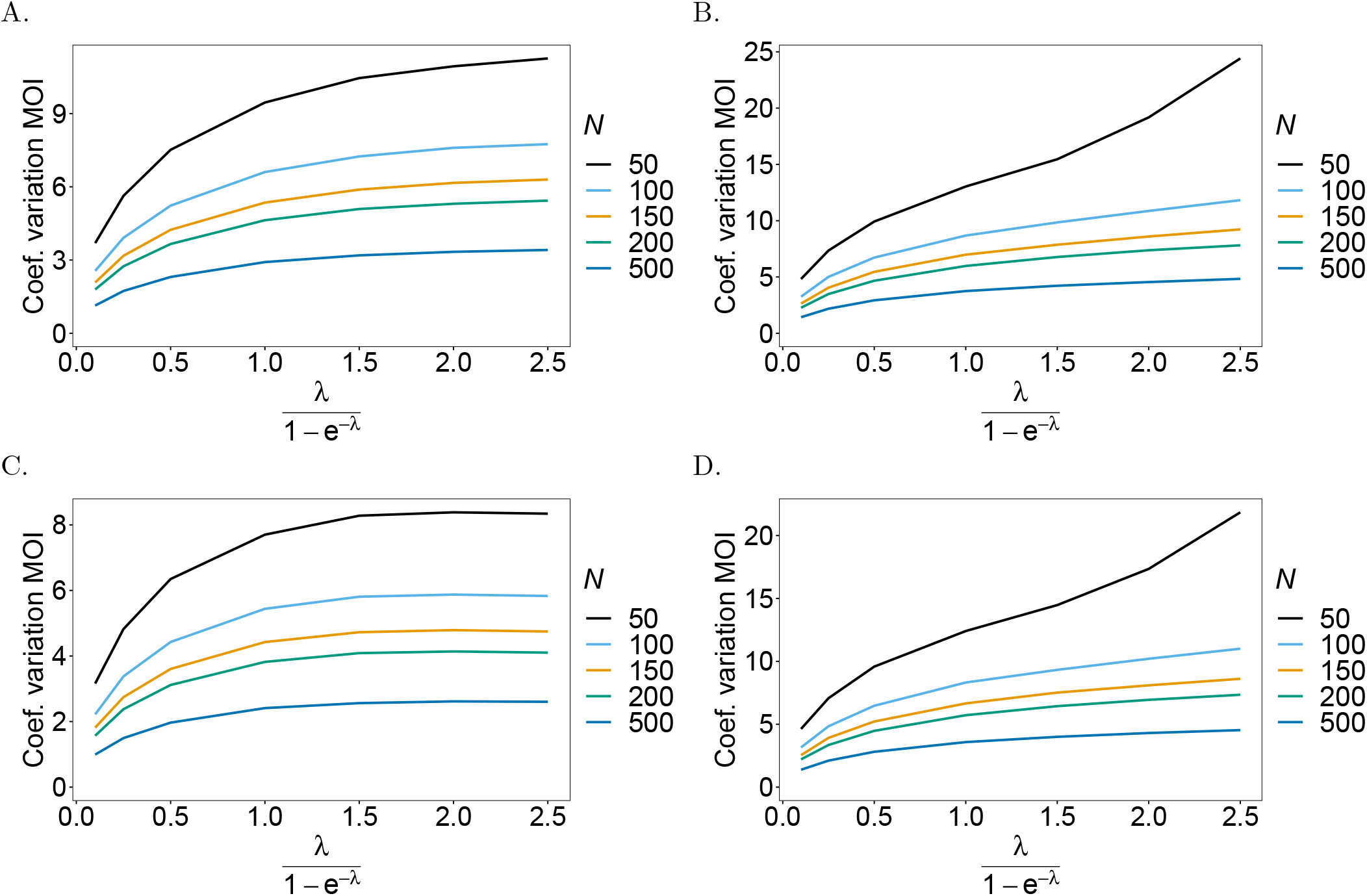
Variance of MOI estimates: Shown is the coefficient of variation (CV) of the mean MOI estimates in % as a function of the true mean MOI (i.e., for a range of MOI parameters). Concerning the genetic architecture *n*_1_ = *n*_2_ = 2 is assumed in (A, C) and *n*_1_ = 4, *n*_2_ = 7 in (B, D). The balanced haplotype frequency distributions of Table 1 are assumed in (A, B) and the unbalanced ones in (C, D). Colors correspond to different sample sizes.

#### Linkage disequilibrium

We estimated the quality of the MLEs as inputs for the estimation of linkage disequilibrium. While we do not have any preference for any of the measures of LD, we focused on three measures, namely, *D′, r*^2^, and *Q*^*\**^ (see eqs.16a, 16b, 16c, respectively). Note that, for *n*_1_ = *n*_2_ = 2, the measures *r*^2^ and *Q*^*\**^ are equivalent. In the case of a balanced true haplotype frequency distribution, all haplotypes are equally distributed in the population. Therefore, the frequencies are in linkage equilibrium (LE), i.e., *D′* = *r*^2^ = *Q*^*\**^ = 0. In general, the estimates of LD obtained from the MLEs are close to their expected values (see Fig 7), however, they cannot be exactly zero, because all LD measures are non-negative. If the frequencies are in LE, in relative terms, the estimates of LD using *D′* depart from LE, more than those based on *r*^2^ and *Q*^*\**^ (see Fig 7). This is explained by the fact that, *D′* depends more on the values of haplotype frequencies than *r*^2^ and *Q*^*\**^ [22]. A comparison between Figs 7A, B, and C, D shows that the error is larger for larger values of *n*_1_ and *n*_2_. This result is a reflection of the larger bias of the estimator due to the increased number of haplotypes for larger *n*_1_ and *n*_2_.

**Fig 7.**
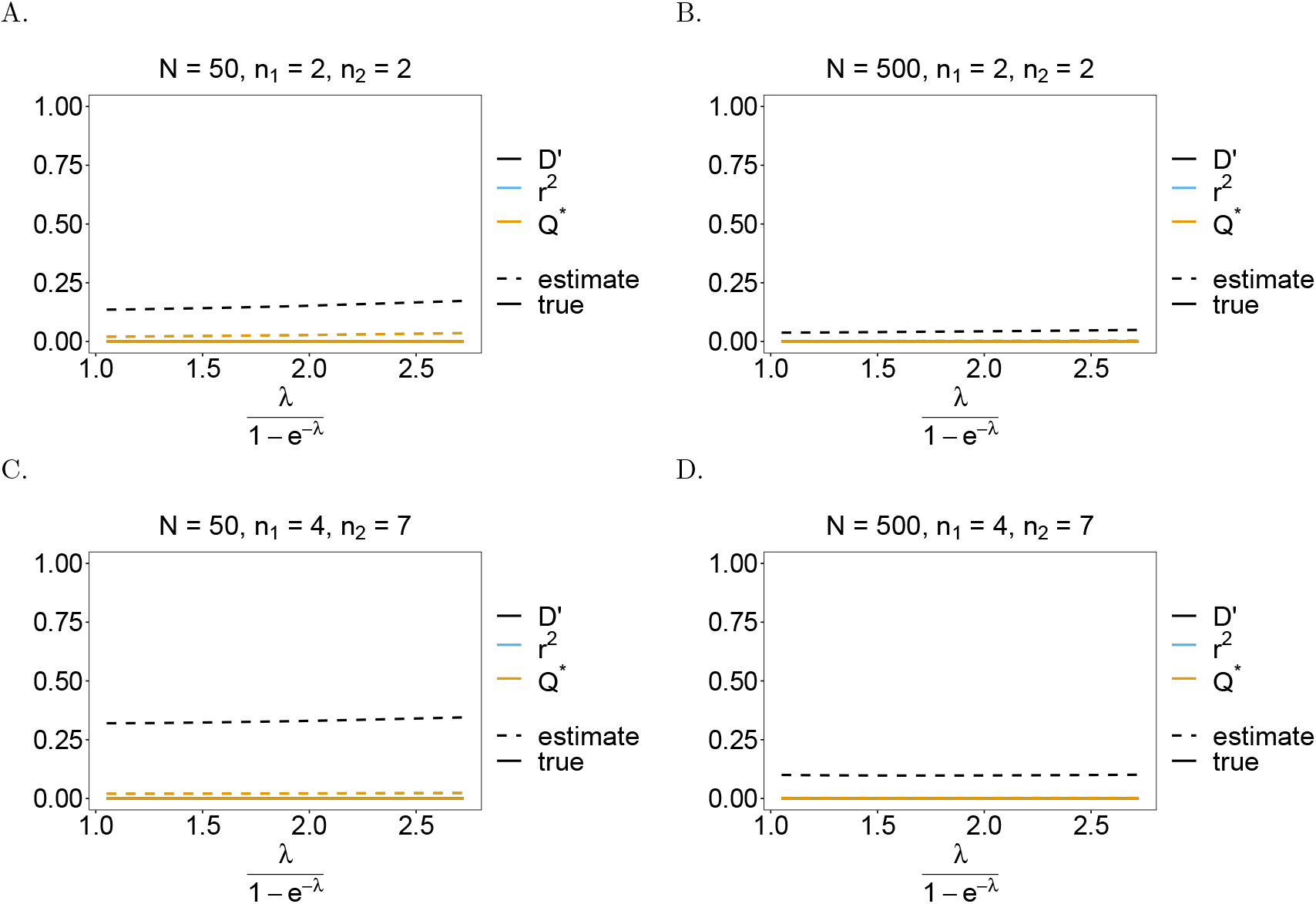
Linkage disequilibrium (LD) – the balanced case: Shown are estimates of LD plotted against the mean MOI. The balanced distributions of Table 1 for the genetic architectures *n*_1_ = *n*_2_ = 2 (A, B) and *n*_1_ = 4, *n*_2_ = 7 (C, D) are assumed. The estimates presented are for small and large sample sizes, i.e., *N* = 50 (A, C) and *N* = 500 (B, D), respectively. Colors correspond to different LD measures. The solid and dashed lines show the true and estimated LD values.

##### Importantly, the error is reduced as sample size increases

Assuming an unbalanced true haplotype frequency distribution, the estimates of LD are close to the true values (see Fig 8). The haplotype distributions are no longer in LE. In this case, the estimates of LD based on *D′* depart less from their true values than for balanced true haplotype frequency distributions (compare Figs 7 and 8). This is because, in the unbalanced distribution, one of the haplotypes is predominant with a high frequency, while the remaining haplotypes are rare. Therefore, the estimated value of *D′* mostly depends on the estimated frequency of the predominant haplotype, which is estimated with low bias (see Fig 3). The deviations from the true value decrease with increasing sample size. The deviations of *r*^2^ and *Q*^*\**^ from their true values are slightly higher for unbalanced frequency distributions, but still smaller than for *D′*.

**Fig 8.**
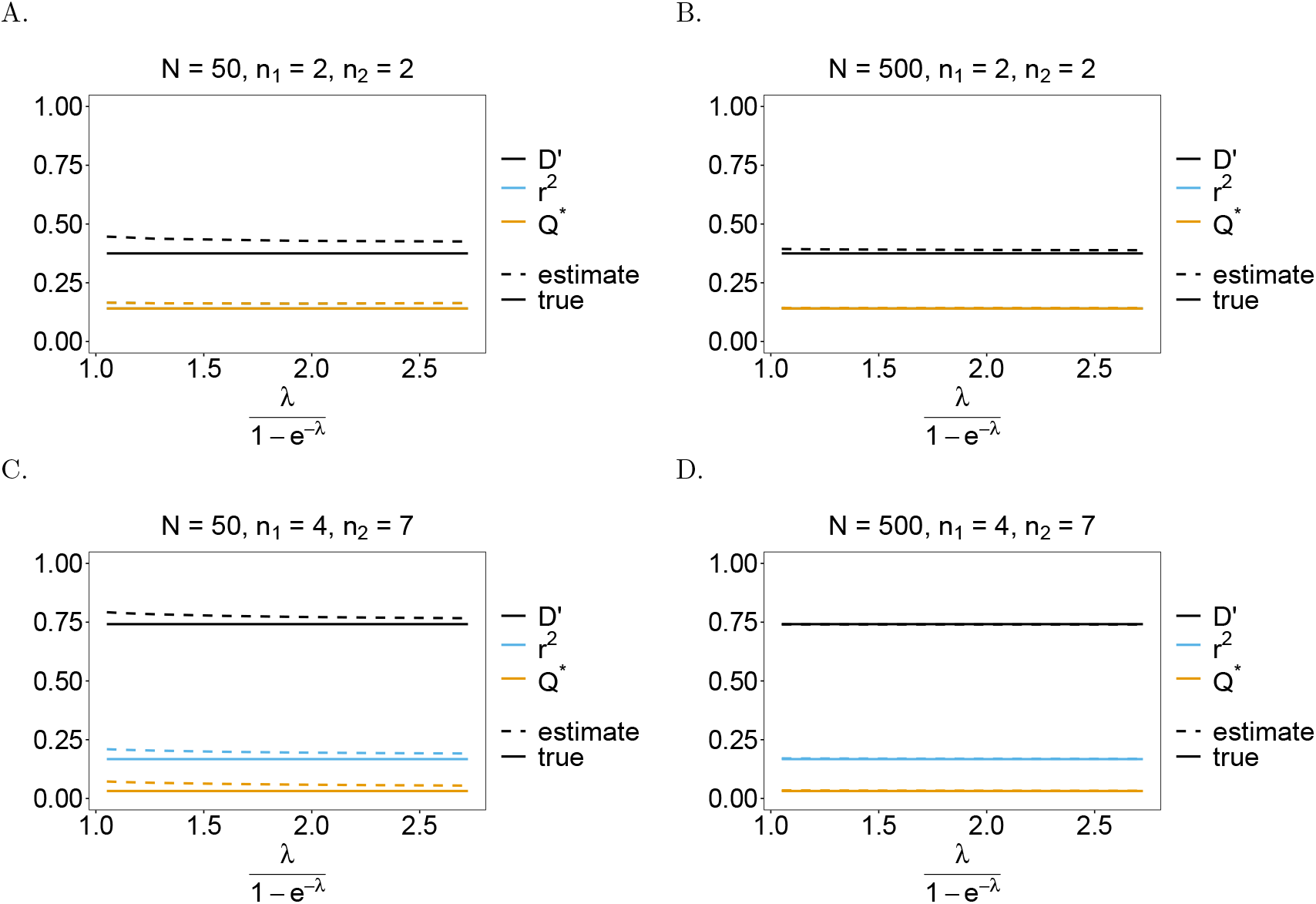
Linkage disequilibrium (LD) – the unbalanced case: Shown are estimates of LD plotted against the mean MOI. We assume the unbalanced haplotype distributions of Table 1 for the genetic architectures *n*_1_ = *n*_2_ = 2 (A, B) and *n*_1_ = 4, *n*_2_ = 7 (C, D). The estimates presented are for small and large sample sizes, i.e., *N* = 50 and *N* = 500, respectively. Colors correspond to different LD measures, while the solid line shows the true value of LD and the dashed line shows the estimates.

## Discussion

Disease surveillance, enhanced by genetic/molecular techniques, became increasingly feasible and popular to monitor infectious diseases [31] such as malaria [16]. Due to considerable efforts and resources devoted to malaria control, the disease burden was reduced substantially in the last two decades [8], with some endemic areas shifting their goals from disease control to elimination. However, there is an upward trend in worldwide malaria incidence and mortality since 2018 and the disease remains highly prevalent in many countries, particularly in Sub-Saharan Africa [8]. Areas with high malaria endemicity harbor substantial genetic variation, e.g., caused by ectopic recombination in the var gene families [32], which needs to be constantly monitored.

Importantly, genetic variation in the pathogen, e.g., msp1 and msp2 in *P. falciparum* can affect disease severity [33]. Furthermore, when aiming for eradication, transmission intensities and routes of transmission have to be monitored closely. Additionally, successful malaria control and eradication attempts are challenged by the emergence and spread of antimalarial-drug resistance and *Pf* HRP2/3 gene deletions [8, 34, 35].

In malaria, molecular surveillance became widely used to monitor genetic diversity [16]. In areas of high transmission, the genetic diversity of pathogen antigens is relevant for the clinical presentation of the disease. Moreover, changes in genetic diversity can reflect the effectiveness of control interventions [36], with decreasing diversity being indicative of successful control measures, and vice versa. Regarding the latter, the increase in genetic diversity in Venezuela between 2004 and 2017, after the economic situation worsened and affected malaria control programs, was obvious and particularly measurable by MOI [37]. In areas of low transmission, molecular surveillance can be informative of routes of transmission, as was described in Colombia [38]. This is particularly important in the context of elimination, to distinguish whether local disease outbreaks are caused by migrational events reflecting deficits in measures to contain outbreaks locally, or from relapses or recrudescence being indicative of deficits in diagnostics or treatments. Diagnostic failures can be the result of the widespread prevalence of *Pf* HRP2/3 deletions, while recrudescence can be a result of widespread drug resistance. Molecular methods are not just adequate to estimate the frequency/prevalence of resistance, e.g., [19, 39], but also to explain and reconstruct the history of drug-resistance evolution [20]. Importantly, the population genetics of *Plasmodium* differs from standard population genetics, due to the organism’s specific transmission cycle [40–42]. More precisely, the processes of selection and recombination cannot be decoupled [43]. This can even result in genome-wide reductions of genetic variation caused by selection in low transmission areas, i.e., in areas with low MOI [15, 43]. Similarly, patterns of linkage disequilibrium (LD) indicative for selection, have to be carefully interpreted in the context of the distribution of MOI. LD is expected to be higher in areas with low malaria endemicity [44], because the effective recombination rates are reduced by low MOI.

To adequately estimate pairwise LD in malaria and similar diseases, we introduced a statistical model to estimate the frequency of pathogenic variants alongside MOI from a pair of multi-allelic marker loci (with *n*_1_ and *n*_2_ alleles segregating at the first and second marker, respectively). This genetic architecture can be interpreted differently. A locus can be an STR (microsatellite) marker (as in the empirical example provided), a SNP which is not necessarily bi-allelic, or a micro haplotype, i.e., a set of DNA sequences or a panel of SNPs in short non-recombining regions, for which phased molecular data can be obtained. The latter is increasingly feasible by third-generation sequencing methods [45]. In this context, MOI follows the definition of [17], i.e., as the number of super-infections during one disease episode. This neglects the possibility of co-transmission of several pathogenic variants at one infectious event (co-infection). The type of molecular data to which the proposed method is applicable does not provide enough resolution to adequately address co-infections. For more discussion and justification of this assumption, see [17].

The proposed method provides maximum-likelihood estimates (MLE) of (two-locus) haplotype frequencies and the distribution of MOI, assuming an underlying Poisson distribution. The latter assumption conveniently reduces MOI to a single parameter. The frequency estimates can be used as plugin estimates to derive LD measures, which have to be interpreted in the context of MOI. The statistical model is too complex to allow a closed-form solution to derive the MLE. Therefore, the expectation-maximization (EM) algorithm was employed, which provides a numerically stable method to derive the MLE. Because the parameter space is constrained, i.e. the frequencies are elements of an (*n*_1_*n*_2_ *−* 1)-dimensional simplex and the MOI parameter is positive, Newton-Raphson methods harbor the problem that the parameter updates might fall out of the admissible parameter space. Specifically, such methods are sensitive to initial parameter choices. The EM-algorithm does not share this problem, but convergence might be slow close to the maximum. The numerical investigations revealed that computational time is sufficiently fast (cf. [46] Chapter 2).

Concerning the genetic architecture, the numbers of segregating alleles *n*_1_ and *n*_2_ are arbitrary. However, the method might suffer from the curse of dimensionality if the numbers of alleles are too large. For instance, for typical sample sizes *N*, up to 30 alleles can be commonly observed at highly variable STR markers. As an example *n*_1_ = *n*_2_ = 30 leads to 900 possible haplotypes (most of which will not even be realized in the population), such that the number of model parameters by far exceeds realistic sample sizes. Notably, in practice, the numerical implementation of the method is not affected by the curse of dimensionality, because typically only a few alleles are observed in a sample at each marker. However, for pathologic examples, implementations of the EM algorithm might exceed the RAM of standard desktop computers.

When creating pairwise LD maps (e.g., Figs 1, 2) between *L* markers, the method needs to be applied 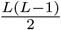 times, thereby yielding 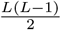 estimates for the MOI parameter and the allele frequencies of each marker can be obtained by marginalization from *L −* 1 of the two-marker estimates. While it is undesirable not to obtain a single estimate for the MOI parameter, it is justified by three reasons. First, if only pairwise LD is of interest, the MOI estimates are of secondary importance. Second, one could average the 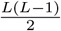 estimates, and use this in a second step as a common plugin estimate for a new estimation of all 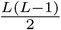 two-marker frequencies, thereby heuristically avoiding the different MOI estimates. Third, a full-haplotype-based method, which derives the frequency distribution for haplotypes characterized by all available markers (and then all two-marker distributions by marginalization) can be undesirable due to the curse of dimensionality, or because of missing data. For the former, assume *L* = 10 markers with 5 alleles segregating at each marker in a sample of size *N* = 200. This results in 9 765 625 possible haplotypes by far exceeding sample size, while there are only 25 possible two-marker haplotypes for each parameter combination, being much smaller than sample size. For the latter, in practice molecular assays sometimes fail, resulting in missing values. In a full-haplotype-based method, only samples can be used, for which molecular assays produced data at all markers. For each 2-marker comparison, more samples can be retained from the data than for a full-haplotype-based method.

The quality of the proposed method was investigated by numerical simulations and showed overall desirable properties. Particularly, bias is low, and the simulation suggests asymptotic unbiasedness. This was explicitly proven for the corresponding method using a single molecular marker [47] and is also in agreement with the corresponding method using a genetic architecture of multiple bi-allelic (SNP) markers [19]. Note, in the case of a single molecular marker, the model falls in the class of exponential families [48], by which the asymptotic properties of the estimator follow. With two or more markers, there is no obvious way to rewrite the model as an exponential family, and it is questionable whether it is possible. Therefore, the asymptotic properties do not immediately follow. However, the results from the single-marker case and the numerical investigations performed here, support that the estimator has desirable small- and large-sample properties.

The quality of LD estimates was assessed through the standard measures, namely, *D′, r*^2^, *Q*^*\**^, and ALD. In general, multi-allelic LD measures are more difficult to interpret than biallelic LD measures [22]. All these measures harbor limitations. Nevertheless, these are standard in evolutionary-genetic analyses and the performance of the proposed method to obtain plugin estimates for LD measures is sufficiently accurate. However, caution is required when choosing an appropriate LD measure.

As an example, we applied the method to an empirical dataset from Cameroon consisting of molecular information at markers flanking the *Pfdhfr* and *Pfdhps* loci in *P. falciparum* malaria, associated with drug resistance. The application confirmed that the method properly captures the intuitively expected behavior, giving further evidence for the appropriateness of the method.

The method is implemented in an easy-to-use R script available as supporting information, on GitHub at https://github.com/Maths-against-Malaria/MultiAllelicBiLociModel (this version will be further maintained) and on Zenodo at https://doi.org/10.5281/zenodo.8289710. For a description and examples of how to use the R script, see S1 User Manual and https://github.com/Maths-against-Malaria/MultiAllelicBiLociModel.

## Methods

The statistical model described here adapts the framework of [19] to estimate MOI and haplotype frequencies to the case of two multialelic loci, e.g., microsatellite markers. Importantly, we define MOI as the number of super-infections, i.e., independent infectious events that occur during the course of the disease (Fig 9), while assuming no co-transmission (co-infections).

**Fig 9.**
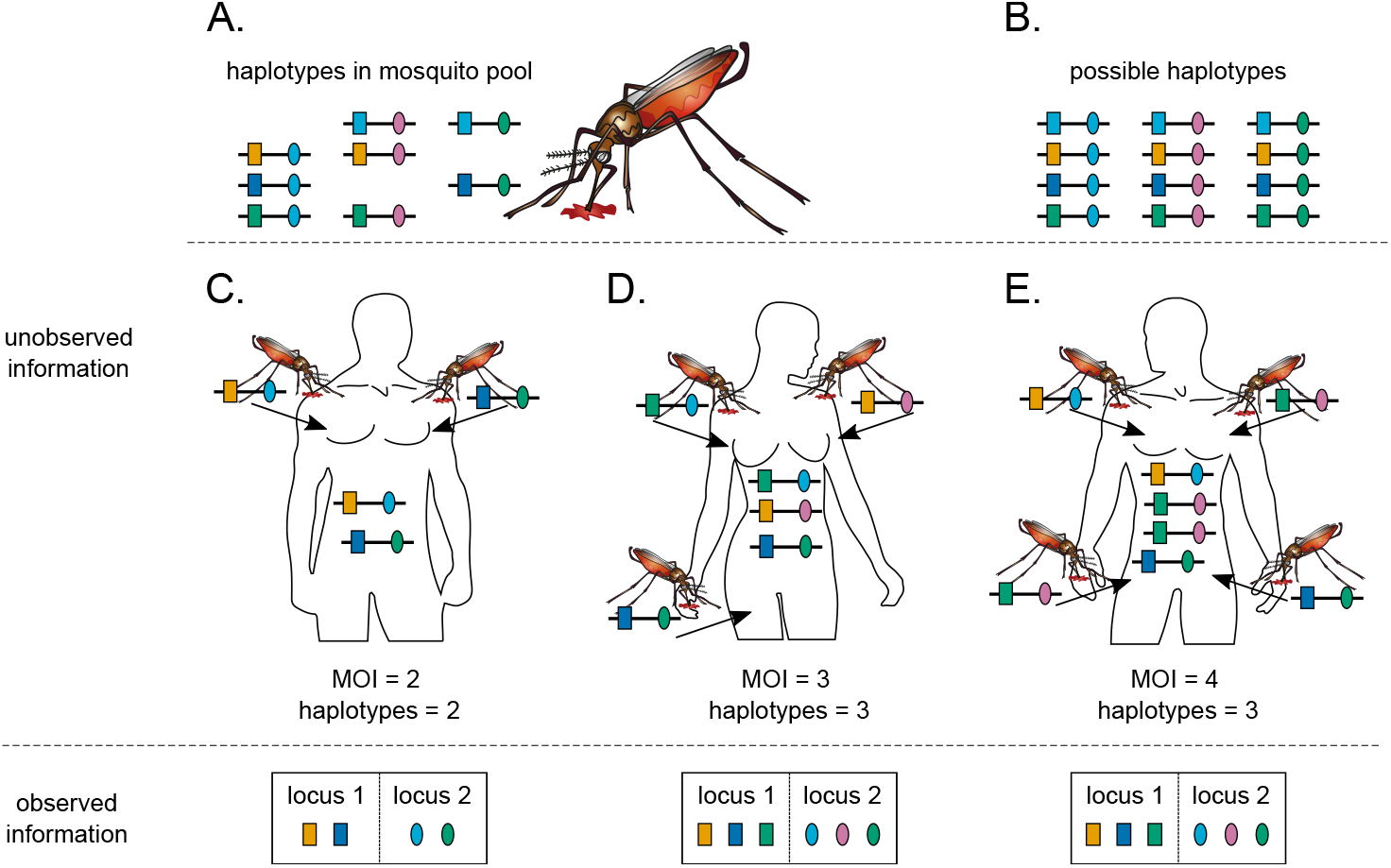
Super-infections and haplotypes phasing: Illustrated are different infective events with haplotypes from the same pathogen population. Panel **(A)** shows the haplotypes present in the parasites population, with each haplotype identified by two loci markers (different shapes), and several alleles at each locus (different colors). Considering the number of loci, and the alleles observed at each locus, there is in theory more haplotype that could be present in mosquito population. All the possible haplotypes are shown in panel **(B)**. Panel **(C)** illustrates an infection with two different infecting haplotypes (i.e., MOI *m* = 2). The corresponding haplotype information (bottom-left) is ambiguous as it is impossible to reconstruct the infecting haplotypes with certainty. Panel **(D)** is similar to Panel **(C)** with three different infecting haplotypes (MOI *m* = 3). The corresponding haplotype information is ambiguous in this case as well (bottom-middle). **(E)** illustrates a super-infection with three different infecting haplotypes, with one of the haplotypes infecting twice (i.e., MOI *m* = 4). The haplotype information in this case is also ambiguous. Note that, in all cases, it is impossible to be confident in identifying how many times the hosts were infected. MOI information is typically ambiguous.

### Statistical model

Let us assume pathogen haplotypes, denoted ***h***, characterized by two loci (markers) with *n*_1_, *n*_2_ alleles, at the first and second locus, respectively. A haplotype is represented by a vector of length two indicating its allelic configuration at each locus, i.e., ***h*** = (*h*_1_, *h*_2_), with *h*_1_ ∈ *{*0, …, *n*_1_ *−* 1*}* and *h*_2_ ∈ *{*0, …, *n*_2_ *−* 1*}*. A total of *H* = *n*_1_*n*_2_ haplotypes are possible with such a genetic architecture. We denote the set of all possible haplotypes by *ℋ*, hence,

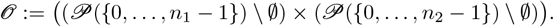

We order the *H* possible haplotypes by the numbers 1, …, *H*, such that haplotype ***h*** corresponds to the number 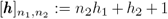. (As an example, for *n*_1_ = 2, *n*_2_ = 4 haplotype (1, 2) corresponds to the number: 4 *·* 1 + 2 + 1 = 4 + 2 + 1 = 7.)

The frequency of haplotype ***h***, denoted by *p*_***h***_, is its relative abundance in the pathogen population. Considering all possible haplotypes, we denote haplotype frequencies with the vector ***p*** := (*p*_***h***_)_***h***∈*ℋ*_ = (*p*_1_, …, *p*_*H*_), where *p*_*k*_ = *p*_***h***_ if 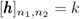. Note that in practice, among the *H* possible haplotypes, only a few are present in the pathogen population, i.e., *p*_*k*_ = 0 for most haplotypes (see Fig 9A,B).

Here, assuming only disease-positive individuals, the number of (super-)infections during the same disease episode, referred to as multiplicity of infections (MOI), follows a conditional (or positive) Poisson distribution [19, 30]. The probability that an individual is (super)-infected exactly *m* times (MOI=*m*) is,

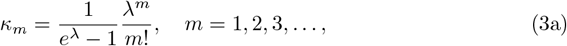

where *λ* is the parameter of the distribution. Moreover, the probability generating function (PGF) of the distribution is

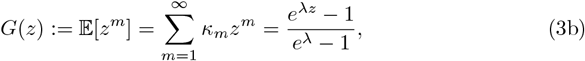

and the mean MOI is (cf. [30])

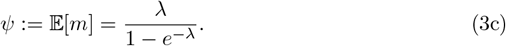

During an infective event, only one haplotype is randomly sampled from the pathogen population and transmitted to the host [19, 30]. Assuming a super-infection with MOI = *m*, an individual is infected *m*_***h***_ times with haplotype ***h*** (see Fig 9C-E). The infection is described by ***m*** := (*m*_***h***_)_***h***∈*ℋ*_ = (*m*_1_, …, *m*_*H*_) such that 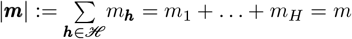. Therefore, given MOI *m*, the probability of infection ***m*** is

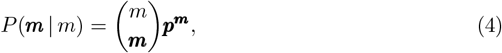

where 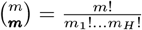, and 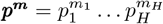.

The infection vector ***m*** and the corresponding MOI *m* are unknown in practice since they cannot be observed from a clinical specimen (see Fig 9C-E). Moreover, due to the lack of phasing, the presence of multiple different haplotypes in a clinical sample yields ambiguous genetic information [19, 49]. Here, we assume that the absence/presence of alleles at considered loci is the only available information.

Given an infection, its allelic information is denoted by the vector ***x*** = (*x*_1_, *x*_2_), where *x*_1_ and *x*_2_ are the set of alleles detected at the first and second locus, respectively. We assume that all alleles present in an infection are detected and there are no erroneous detection. The allelic information *x*_1_, is a subset of all alleles detectable at the first locus. Therefore, *x*_1_ is an element of the powerset of *{*0, …, *n*_1_ *−* 1*}*, i.e., *x*_1_ ∈ *𝒫*(*{*0, …, *n*_1_ *−* 1*}*) \ ∅. Since only disease-positive samples are considered, the empty set is excluded. Equivalently, *x*_2_ ∈ *𝒫*(*{*0, …, *n*_2_ *−* 1*}*) \ ∅. Therefore, the set of all possible observations is given by the following cartesian product

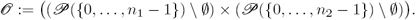

There is a total of 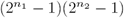 possible observations. Note that a particular observation ***x*** can result from different infections ***m***, i.e., ***m*** → ***x*** (cf. Fig 9D-E). Given MOI *m* and observation ***x***, the set of all such infections is denoted by

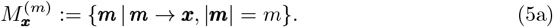

Furthermore, for an observation ***x***, we define the set of all haplotypes that could potentially be present in ***x*** as

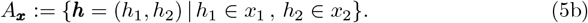

The set of all observations with at most the same alleles detected at each locus as in ***x*** is denoted by

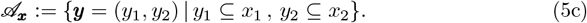

We define the partial order “≼” on the set of all possible observations, such that ***y*** ≼ ***x*** is equivalent to ***y*** ∈ *𝒜*_***x***_. If ***y*** ≼ ***x*** and ***x*** ≠ ***y*** we write ***y*** ≺ ***x***. Therefore, we can express 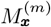 as

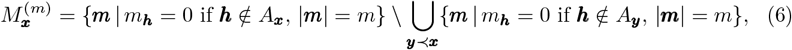

*A*_***y***_ is defined as in (5b) for observation ***y*** [19]. Consider a host infected *m* times (MOI= *m*), the probability that the actual infection is ***x***, is given by

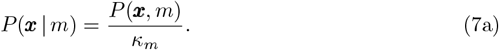

Hence,

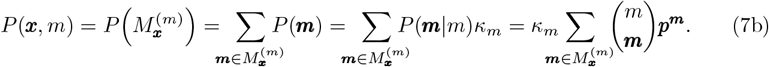

The probability of observation ***x*** becomes

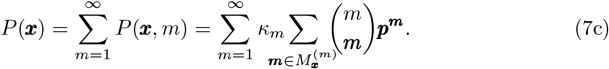

The simplified notation *P*_***x***_ will be used from now on, in place of *P* (***x***) to denote the probability of observing ***x***. Furthermore, let |***x***| denote the cardinality of the set ***x***. By using (5a) and the inclusion-exclusion principle, the inner sum on the right-hand side of (7c) can be rewritten [19] as

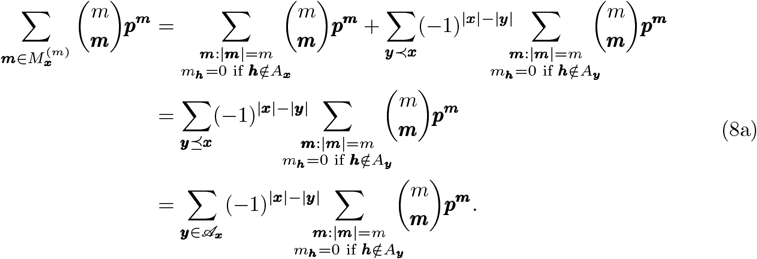

Therefore, the probability of observing ***x*** in (7c) becomes

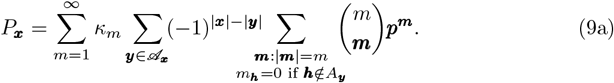

By the multinomial theorem, we have

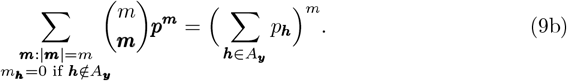

By changing the order of summation and using the PGF (9a) becomes

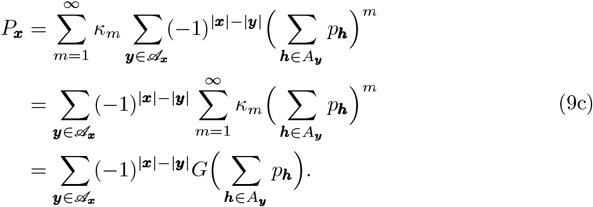

Note, the above equation is correct (cf. eq. 9c) for any distribution of MOI. If we impose the condition Poisson distribution, the MOI parameter *λ* occurs in the PGF. Therefore, the probability *P*_***x***_ depends on the MOI parameter *λ* and the haplotype frequencies ***p***. The parameter space of the conditional Poisson model is

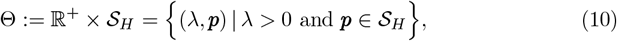

where, 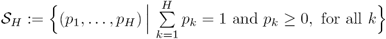 is the *H −* 1-dimensional simplex.

The true parameters, here subsumed by the vector ***θ*** = (*λ*, ***p***), are unknown and can be inferred from empirical data. Assume a dataset *𝒳* consisting of *N* observations ***x***^(1)^, …, ***x***^(*N*)^, where the notation 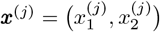 is used for the *j*th observation. For the dataset *𝒳*, let *n*_***x***_ be the number of times observation ***x*** is made. Naturally,

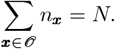

Using (9c), the likelihood function of the parameter ***θ*** = (*λ*, ***p***) given the data *𝒳* is given by

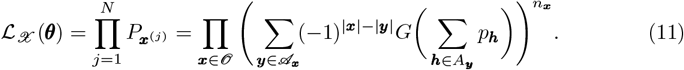

Hence, the log-likelihood function becomes

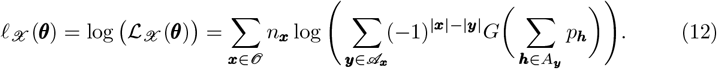

To obtain the maximum-likelihood estimate (MLE) 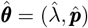 the log-likelihood function needs to be maximized. The complexity of the log-likelihood function does not permit a closed solution, and must be maximized numerically. For this purpose the expectation-maximization (EM)-algorithm will be used [50]. This will be discussed in The maximum-likelihood estimate.

Note that (12) holds for any distribution of MOI. If the positive Poisson distribution is replaced, only the PGF *G* and the parameter ***θ*** need to be modified.

#### Assessing bias and variance of the estimator

MLEs usually have desirable asymptotic (large sample size) properties. In practice, sample size is often limited, and the quality of the estimator needs to be investigated for small and intermediate sample sizes. Because no explicit solution exists for the MLE, its performance in terms of bias and variance needs to be investigated by numerical simulations. Here, we adapt the approach of [19, 47, 51].

Bias and variance of the MLE will be affected by: (i) sample size *N* ; (ii) the genetic architecture, i.e., the number of alleles *n*_1_, *n*_2_ segregating at each locus; (iii) the value of the MOI parameter *λ*; (iv) the frequency distribution of haplotypes ***p***.

To investigate the properties of the MLE for a representative range of parameters we proceeded as follows (parameters used in the simulation study are described below and summarized in Table 1). For a set of parameters (*N, n*_1_, *n*_2_, *λ*, ***p***) we generated *K* = 100 000 datasets *𝒳*_1_, …, *𝒳*_*K*_ of size *N* according to the model (9c). For each dataset *𝒳*_*k*_, the MLE 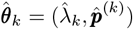 was calculated. From each 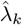 the mean MOI 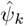 was calculated according to (3c). The bias and variance of the mean MOI *ψ* were estimated as

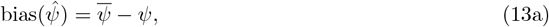

and

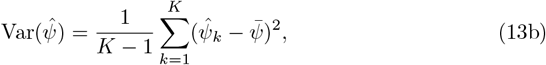

where

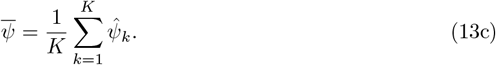

To allow comparisons between different parameter ranges it is more appropriate to consider the relative bias and coefficient of variation, which are independent of the scale, i.e.,

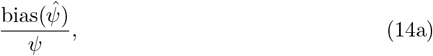

and

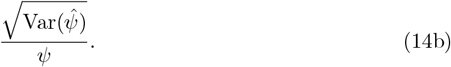

For each haplotype frequency *p*_***h***_, bias and variance were defined in the same way with obvious modifications.

**Table 1.**
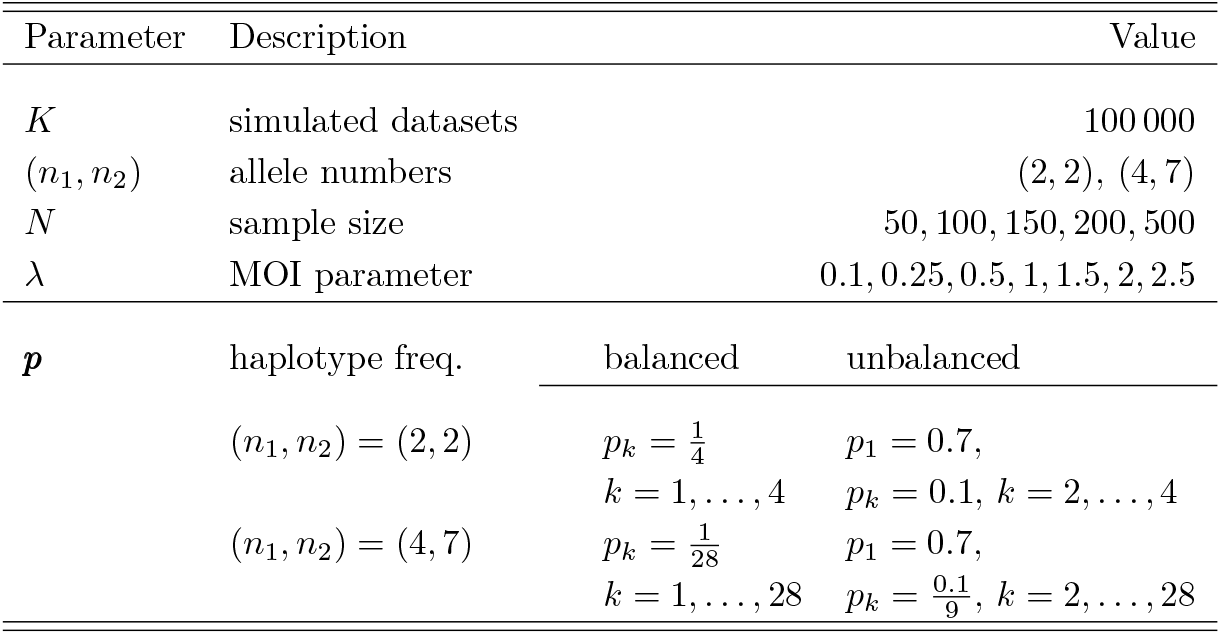
Summary of model parameters chosen for simulations in the assessment of the performance of the estimator.

##### Genetic architecture

Considering the number of alleles at each of the 2 loci considered, to investigate the performance of the estimator, we assumed the cases, (i) *n*_1_ = *n*_2_ = 2 and (ii) *n*_1_ = 4, *n*_2_ = 7. This yields, respectively, 4 and 28 possible haplotypes in total. The simulation is based on those cases, as an exhaustive simulation on all possible configurations would be too expensive computationally due to the curse of dimensionality.

The model is not restricted to the number of alleles per locus. However, ∼ 30 alleles per locus would result in ∼ 900 parameters, which would require a sample size of *N >* 1 000 to obtain reliable results.

#### MOI parameter

Concerning the MOI parameter we chose *λ* = 0.1, 0.25, 0.5, 1, 1.5, 2, 2.5, corresponding to a mean MOI *ψ* = 1.05, 1.13, 1.27, 1.58, 1.93, 2.31, 2.72. In the case of malaria, this corresponds to low transmission *ψ <* 1.27, intermediate transmission 1.27 ≤ *ψ <* 1.93, and high transmission *ψ* ≥ 1.93 [51].

#### Haplotype frequency distribution

The following haplotype frequency distributions ***p*** were chosen. First, a completely uniform (balanced) distribution was chosen, i.e., each of the *H* = *n*_1_*n*_2_ haplotype had the same frequency,

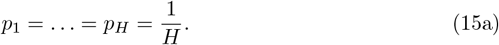

Then, an unbalanced distribution with one predominant haplotype was chosen. The frequency of the predominant haplotype was chosen to be 70%, while the remaining haplotypes all had the same frequency. In particular, we chose

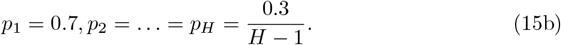

For the genetic architecture *n*_1_ = *n*_2_ = 2 this yielded, *p*_1_ = 0.7, *p*_2_ = *p*_3_ = *p*_4_ = 0.1 and *n*_1_ = 4, *n*_2_ = 7 yielded *p*_1_ = 0.7, *p*_2_ = … = *p*_28_ = 0.011.

#### Sample size

The performance of an estimator is usually affected by sample size. We investigate the effect of sample size in our numerical investigations by constructing datasets of size *N* = 50, 100, 150, 200, 500, which are typical sample sizes ranging from areas of low to high transmission in diseases like malaria.

The simulation study as well as the graphical outputs are implemented in R [52].

The code is available at https://github.com/Maths-against-Malaria/MultiAllelicBiLociModel and on Zenodo at https://doi.org/10.5281/zenodo.8289710.

### Linkage disequilibrium

Linkage disequilibrium (LD) also known as gametic disequilibrium is a measure of the association between alleles at different loci, i.e. statistical dependence. LD is calculated from haplotype frequency estimates. Two loci are said to be in linkage equilibrium if the haplotype frequencies coincide with the products of allele frequencies, i.e., statistical independence, otherwise, they are said to be in LD. For two bi-allelic loci, LD measures have rather straightforward interpretations (cf. [53]). For multi-allelic loci LD measures and their interpretation are less straightforward. There exist a variety of multi-allelic LD measures in the literature, each having its advantages and limitations (cf. [22]). Here, to assess the quality of the MLEs when used as plug in estimates to derive LD, we focus on three commonly used measures, i.e., *D′, r*^2^, and *Q*^*\**^.

We further exemplify the three LD measures with an empirical data application. In the data application, we also derive the more recently developed asymmetric conditional LD (ALD) measure introduced in [23], which accounts for asymmetry in the number of alleles observed at each locus.

The various LD measures are defined as follows. Let *x*_*i*_ (0 *< x*_*i*_ *<* 1) denote the frequency of allele *A*_*i*_ (*i* = 1, …, *n*_1_) at the first and *y*_*j*_ (0 *< y*_*j*_ *<* 1) of allele *B*_*j*_ (*j* = 1, …, *n*_2_) at the second locus. The loci are referred to as locus *A* and *B*, respectively. All LD measures are undefined if at least one of the loci is monomorphic.

The measure *D′* defined in [22] is calculated as

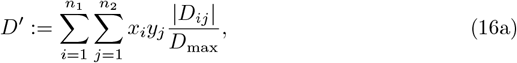

where *D*_*ij*_ = *p*_*ij*_ *− x*_*i*_*y*_*j*_ is the difference between the observed frequency of haplotype *A*_*i*_*B*_*j*_ and the expected frequency *x*_*i*_*y*_*j*_ of *A*_*i*_*B*_*j*_ assuming a random association of the alleles *A*_*i*_ and *B*_*j*_, and

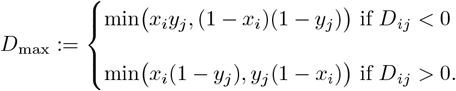

The measure *r*^2^ also known as *D*^*\**^ [22] is defined by

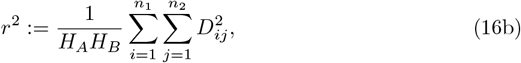

where 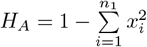 and 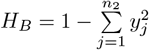 are the Hardy-Weinberg heterozygosities at loci *A* and *B*, respectively. Finally, the measure *Q*^*\**^ is defined in [22] as

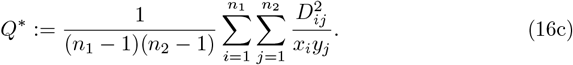

Note, *r*^2^ and *Q*^*\**^ are equivalent for bi-allelic loci, i.e., *n*_1_ = *n*_2_ = 2.

Moreover, [23] defines ALD between the two loci conditioned on *A* by

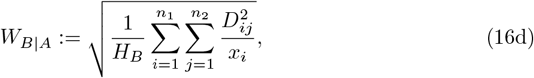

while ALD between loci *A* and *B* conditioned on *B* is defined by

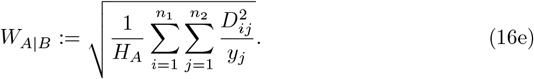

For bi-allelic loci (16d) and (16e) equal and coincide with the square root of *r*^2^, i.e.,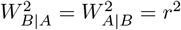.

## Acknowledgements

The molecular dataset used for data applications was provided by Dr. Andrea M. McCollum and Dr. Venkatachalam Udhayakumar, which is gratefully acknowledged.

## Supporting information

**S1 Fig. Linkage disequilibrium at SNP markers:** Shown are estimates of pairwise LD for six SNP marker loci associated with SP drug-resistance, i.e., three codons (51, 59, 108) at *Pfdhfr* and three codons (436, 437, 613) at *Pfdhps*. Panels (A, B) show *r*^2^ values, whereas (C, D) show *Q*^*\**^ values. Furthermore, panels (A, C) correspond to the years 2001/2002, while panels (B, D) correspond to the years 2004/2005. The codons on *Pfdhfr* and *Pfdhps* are highlighted on the maps by horizontal black lines, with the name of the gene and codons are specified on top and below the line, respectively. The thick black lines in panels (A, B) group pairwise LD within each gene. The numbers indicate LD values.

**S2 Fig. Linkage disequilibrium at microsatellite markers:** As in S1 Fig but for a set of genes at two neutral chromosomes, i.e., chromosomes 2 and 3, genes around *Pfdhfr* on chromosome 4, and genes around *Pfdhps* on chromosome 8.

**S1 Mathematical appendix. Mathematical background**.

**S1 User Manual. Description of the usage of the method’s implementation**.

**S1 R Script and example datasets. Zip file containing an R script containing the implementation of the method and a few example datasets**.

